# Identification of Key Proteins Associated with Myocardial Infarction using Bioinformatics and Systems Biology

**DOI:** 10.1101/308544

**Authors:** Ishtiaque Ahammad

**Affiliations:** Department of Biochemistry and Microbiology, North South University, Dhaka, Bangladesh.

**Keywords:** Myocardial Infarction, Heart Attack, Gene Set Enrichment Analysis, Protein-Protein Interaction Network, Bioinformatics, Systems Biology, *In Silico* study

## Abstract

Myocardial infarction, more commonly known as heart attack, is a huge health problem around the world. It is a result of inadequate blood supply to certain parts of the heart and death of heart muscle cells in that region. Although it has been around for a long time, newer and newer ways of probing myocardial infarction is being followed. Bioinformatics and Systems Biology are relatively recent fields to try to give new insights into myocardial infarction. Following the footsteps of others, this *in silico* study has tried to chime in on the investigation into myocardial infarction. The study began with the gene expression omnibus dataset uploaded to the NCBI from a whole-genome gene expression analysis carried out at Mayo Clinic in Rochester, Minnesota. From the dataset, differentially expressed genes following first-time myocardial infarction were identified and classified into up-regulated and down-regulated ones. Gene Set Enrichment Analysis (GSEA) was carried out on the up-regulated genes which were statistically significant and corresponded with the NCBI generated annotations. Protein-Protein Interaction Network for these genes was constructed. GSEA revealed 5 transcription factors, 5 microRNAs and 3 pathways significantly associated with them. From the Protein-Protein Interaction Network, 6 key proteins (hub nodes) have been identified. These 6 proteins may open a new window of opportunity for the discovery/design of new drugs for mitigating the damage caused by myocardial infarction.

## Introduction

Myocardial Infarction, also known as heart attack involves the death of cells in the myocardium. It is caused by an unstable ischemic syndrome (Thygesen *et al*., 2012). Three universal definitions of myocardial infarction have been given so far.

The first universal definition of myocardial infarction was devised by the First Global MI Task Force based on the elevation and decline of biochemical markers characteristic of myocardial necrosis (Alpert *et al*., 2000).

The Second Global MI Task Force defined myocardial infarction as myocardial necrosis consistent with myocardial ischaemia (Thygesen, Alpert and White, 2007).

The third universal definition of myocardial infarction was developed by a joint taskforce combining four societies: the American College of Cardiology (ACC), the American Heart Association (AHA), the European Society of Cardiology (ESC), and the World Heart Federation (WHF). They classified myocardial infarction into 5 categories according to the myocardial lesion mechanism (Thygesen *et al*., 2012).

Investigation into the molecular mechanisms of myocardial infarction has been in progress for quite some time now. High-throughput SNP analysis, genome-wide scanning, enriched pedigree analysis and many other molecular techniques have been applied (Jefferson and Topol, 2005). Other studies have focused on the cardiac remodeling after an acute myocardial infarction (Vilahur *et al*., 2011; Schirone *et al*., 2017). The role of microRNAs in myocardial infarction and the possibilities of using them in diagnosis and therapy has also been discussed more recently (Sun *et al*., 2017).

Using Bioinformatics to investigate myocardial infarction is also a relatively recent phenomenon (Gao *et al*., 2016). Some of the studies have focused on the differential gene expression during different time periods after a myocardial infarction (Zhang *et al*., 2014) while some of them focused on the miRNA-transcription factor co-regulatory networks during the progression of myocardial infarction (Shi *et al*., 2016).

In this study, differentially expressed genes after myocardial infarction has been analyzed through various tools of Bioinformatics and Systems Biology to shed more light into the molecular mechanisms and key players involved and identify potential targets for drug development.

## Materials and Methods

### Retrieval of Microarray Data

Gene Expression Omnibus Datasets (GEO datasets) from the NCBI website were searched with the term “Myocardial Infarction”. The dataset with the reference series no- GSE48060 was selected for analysis. The expression profiling by array was carried out by Health Sciences Research Department of Mayo Clinic, Rochester, Minnesota. In that study blood samples were collected from 21 control and 31 myocardial infarction patients. Out of the 31 patients, 5 had recurrent events (Suresh *et al*., 2014).

For the present study, those samples were classified into two groups- Myocardial infarction patients without recurrent events (26 samples) and control (21 samples). The purpose of having myocardial infarction as the sole variable was to determine the differentially expressed genes in myocardial infarction.

### GEO2R Analysis

For identifying differentially expressed genes, the samples were subjected to GEO2R analysis (Smyth, 2004) using GEOquery (Davis and Meltzer, 2007) and limma R (Smyth, no date) package. As a result, the most highly differentially expressed 250 genes have been identified. The Benjamini & Hochberg (false discovery rate) method (Benjamini and Hochberg, 1995) was used to adjust the P values. Log2 transformation was applied on the sample data using the auto-detect option in GEO2R.

### Gene Set Enrichment Analysis (GSEA) for the Up-regulated Genes

For GSEA, the gene set enrichment analysis web server, Enrichr (Chen *et al*., 2013; Kuleshov *et al*., 2016), developed by the Ma’ayan Lab was used. For the significantly up-regulated genes, the ARCHS4 (Lachmann *et al*., 2018), miRTarBase 2017 (Chou *et al*., 2018) and BioCarta 2016 (Rouillard *et al*., 2016) databases were used to determine the Transcription Factors, MiRNAs and Pathways associated with them respectively.

### Protein-Protein Interaction Network (PPI Network) Analysis for the Up-regulated Genes

A PPI network for the up-regulated genes were constructed, visualized and analyzed using Cytoscape 3.6.0 (Saito *et al*., 2012). Interactions from 5 databases were imported and merged for constructing the PPI network- iRefIndex (Razick, Magklaras and Donaldson, 2008), APID Interactomes (Alonso-López *et al*., 2016), menthe (Calderone, Castagnoli and Cesareni, 2013), IntAct (Kerrien *et al*., 2012), InnateDB-all (Breuer *et al*., 2013). From the initial network, connected components were identified and the largest individual connected component was extracted to generate a separate second network. The second network containing the largest individual connected component was visualized using the yFiles organic layout in Cytoscape. The node size and colour were mapped based on the average shortest path length between the nodes.

CytoNCA plug-in (Li *et al*., 2017) within Cytoscape was used to identify the “hub nodes” which are crucial proteins in a PPI network (He and Zhang, no date). Hub nodes were defined in terms of degree centrality (DC), betweenness centrality (BC), and closeness centrality (CC) and the parameter was set to “without weight”.

## Results

### Identification of differentially expressed genes

- Out of the 250 differentially expressed genes identified by GEO2R analysis, the 70 genes met the criteria for statistical significance of Adjusted P value of <0.05.
- Out of the 70 significantly differentially expressed genes, 32 were up-regulated and 38 were down-regulated.
- Out of the 32 up-regulated genes, 11 corresponded with NCBI generated annotation.
- Out of the 38 down-regulated genes, 29 corresponded with NCBI generated annotation.
- These 11 up-regulated genes listed below were selected for further analysis (GSEA and PPI network analysis).

**Table 1:** Significantly up-regulated genes that corresponded with NCBI generated annotation in GEO2R

| ID | Adjusted P Value | P Value | logFC | Gene Symbol |
| --- | --- | --- | --- | --- |
| 213350_at | 0.00272 | 9.95E-08 | -0.692 | RPS11 |
| 237444_at | 0.009 | 2.03E-06 | -0.64 | KIF13A |
| 211074_at | 0.03394 | 2.98E-05 | -0.58 | FOLR1 |
| 244889_at | 0.009 | 1.89E-06 | -0.562 | LOC400499///LOC388210 |
| 224846_at | 0.03385 | 2.84E-05 | -0.44 | SHKBP1 |
| 223925_s_at | 0.03847 | 3.59E-05 | -0.414 | MTPN |
| 1566342_at | 0.04686 | 5.50E-05 | -0.409 | SOD2 |
| 208052_x_at | 0.00272 | 9.49E-08 | -0.378 | CEACAM3 |
| 1554360_at | 0.01382 | 4.84E-06 | -0.321 | FCHSD2 |
| 201605_x_at | 0.03065 | 2.27E-05 | -0.311 | CNN2 |
| 227175_at | 0.03065 | 2.26E-05 | -0.236 | MCL1 |

### Gene Set Enrichment Analysis (GSEA) of the Up-regulated Genes

**Table 2:** Top results of Gene Set Enrichment Analysis (GSEA) of the up-regulated genes

| ARCHS4 TFs Coexp |  |  |  |  |  |
| --- | --- | --- | --- | --- | --- |
| Index | Name | P-value | Adjusted p-value | Z-score | Combined score |
| 1 | KLF2_human_tf_ARCHS4_coexpression | 1.49E-05 | 0.002238 | -1.67 | 18.55 |
| 2 | TRERF1_human_tf_ARCHS4_coexpression | 1.49E-05 | 0.002238 | -1.59 | 17.7 |
| 3 | GLI1_human_tf_ARCHS4_coexpression | 0.000499 | 0.005781 | -1.72 | 13.11 |
| 4 | DHX34_human_tf_ARCHS4_coexpression | 0.000499 | 0.005781 | -1.69 | 12.82 |
| 5 | ZNF746_human_tf_ARCHS4_coexpression | 0.000499 | 0.005781 | -1.68 | 12.8 |
| miRTarBase 2017 |  |  |  |  |  |
| Index | Name | P-value | Adjusted p-value | Z-score | Combined score |
| 1 | hsa-miR-7159-3p | 1.03E-07 | 5.47E-05 | -2.2 | 35.46 |
| 2 | hsa-miR-8485 | 0.01054 | 0.09842 | -6.22 | 28.32 |
| 3 | hsa-miR-124-3p | 0.04006 | 0.1163 | -8.56 | 27.54 |
| 4 | hsa-miR-329-3p | 0.002718 | 0.0482 | -4.37 | 25.8 |
| 5 | hsa-miR-362-3p | 0.00269 | 0.0482 | -4.33 | 25.64 |
| BioCarta 2016 |  |  |  |  |  |
| Index | Name | P-value | Adjusted p-value | Z-score | Combined score |
| 1 | The IGF-1 Receptor and Longevity_Homo sapiens_h_longevityPathway | 0.008767 | 0.008767 | -1.52 | 7.21 |
| 2 | Erythropoietin mediated neuroprotection through NF-kB_Homo sapiens_h_eponfkbPathway | 0.007129 | 0.008767 | -1.3 | 6.43 |
| 3 | Cardiac Protection Against ROS_Homo sapiens_h_flumazenilPathway | 0.006035 | 0.008767 | -1.23 | 6.27 |

### PPI Network for the Up-regulated Genes

**Figure 1:**
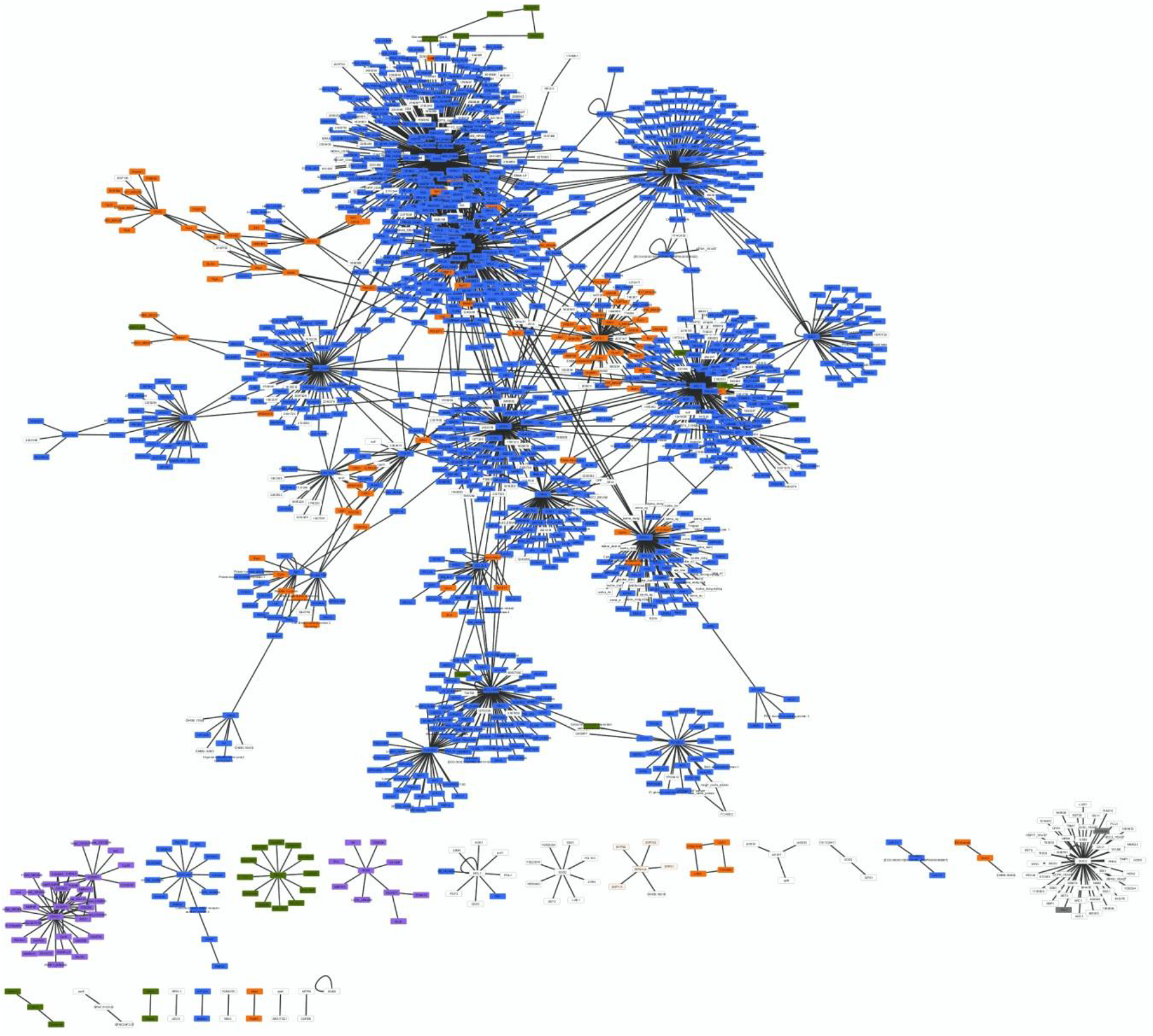
Protein-protein interaction network of the 11 up-regulated genes generated by merging networks from 5 interactions databases. It contains 1519 nodes (proteins), 1830 edges (interactions) and 24 individual connected components. The largest individual connected component has 1333 nodes and the smallest one has only 1 node.

**Figure 2:**
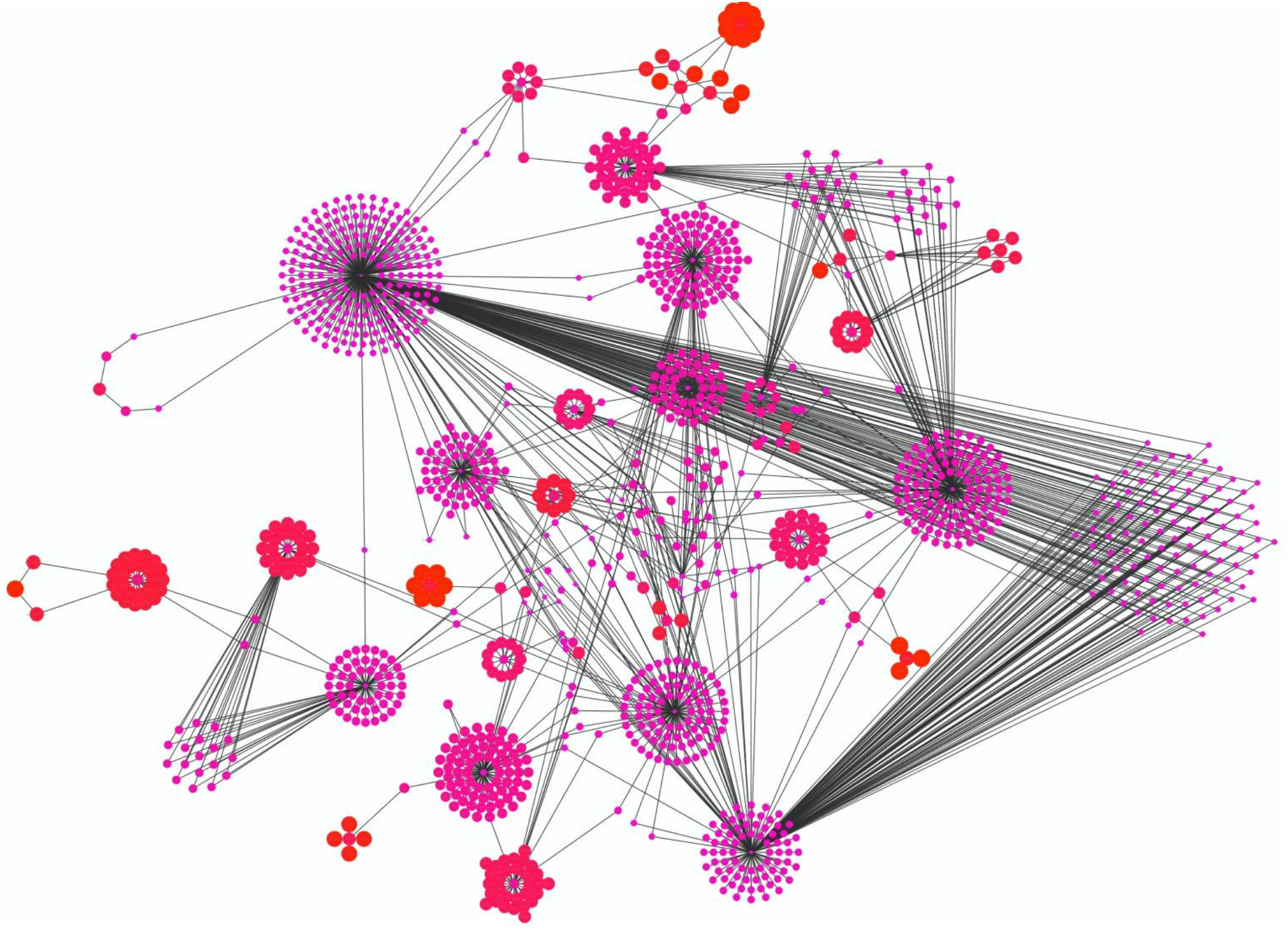
Largest individual connected component (yFiles organic layout) of the initial PPI network of the 11 up-regulated genes. It contains 1333 nodes (proteins) and 1631 edges (interactions). The node sizes and colors are mapped based on the average shortest path length between the nodes. Values range from Pink (2.55) to red (7.50).

### Key Proteins (Hub Nodes) of the PPI Network

**Table 3:** Hub nodes of the largest connected component of the PPI network (based on DC, BC, and CC)

| Degree Centrality |  | Betweenness Centrality |  | Closeness Centrality |  |
| --- | --- | --- | --- | --- | --- |
| Gene | Score | Gene | Score | Gene | Score |
| RPS11 | 350 | RPS11 | 740999.2 | UBC | 0.39350075 |
| RPS11 | 210 | MCL1 | 476977.75 | RPS11 | 0.3615635 |
| MCL1 | 159 | UBC | 409532.62 | RPS11 | 0.3306852 |
| SOD2 | 113 | RPS11 | 408625.16 | TRAF6 | 0.32913268 |
| MTPN | 112 | MTPN | 329884.1 | MTPN | 0.3193479 |

## Disscussion

From the differential gene expression analysis, 11 genes that correspond to NCBI generated annotation were found to be up-regulated (Table 1). Through Gene Set Enrichment Analysis of these 11 genes, several of their molecular mechanisms have been identified. For example, KLF2, TRERF1, GLI1, DHX34, ZNF746 have been found to be the transcription factors influencing the transcription of the gene set (Table 2). From the miRTarBase, 5 microRNAs namely, hsa-miR-7159-3p, hsa-miR-8485, hsa-miR-124-3p, hsa-miR-329-3p, hsa-miR-362-3p have been found to be associated with the up-regulated gene set (Table 2). Using BioCarta pathway database, three pathways have been identified as being associated with the gene set (Table 2). The PPI network of the 11 up-regulated genes contained 1519 proteins and 1830 interactions (Figure 1). From the 24 connected components of the PPI network, the largest one containing 1333 proteins and 1631 edges was extracted, visualized and analyzed as a second network (Figure 2). From the second network, 6 key proteins have been identified as hub nodes. They are RPS11, MCL1, MTPN, SOD2, UBC, and TRAF6 (Table 3).

RPS11, also known as the ribosomal protein S11, is part of the 40S ribosomal subunit (Feo, Davies and Fried, 1992; Kenmochi *et al*., 1998).

MCL1 mainly plays a role in inhibition of apoptosis but two of its isoforms do the opposite (Fung *et al*., 1976). It has been found that MCL1 is crucial for maintenance of homeostasis in myocardium and the decline of MCL1 in the heart is associated with heart failure (Thomas and Gustafsson, 2013). It has been found to be overexpressed in the early stages of myocardial infarction (Matsushita *et al*., 1999).

MPTN, also known as Myotropin, is associated with cardiac hypertrophy (Das *et al*., 2010). It has been considered as a strong predictor of major adverse cardiac events (Khan *et al*., 2007).

SOD2, also known as Superoxide Dismutase 2, is another one of the proteins identified as a hub protein in this study. A recent study has provided insight into the regulation of SOD2 by MicroRNAs as new circulating biomarkers of heart failure (Dubois-Deruy *et al*., 2017).

UBC, also known of Ubiquitin C, is a precursor of polyubiquitin (Board *et al*., 1992). It is one of the sources of ubiquitin (Kimura and Tanaka, 2010). The Ubiquitin proteasome system (UPS) plays an important role in protein degradation and is related with the contraction of the heart (Willis *et al*., 2014).

TRAF6, also known as TNF Receptor Associated Factor 6 or E3 Ubiquitin-Protein Ligase TRAF6, is involved in the mitigation of acute myocardial infarction damage (SUN, HUANG and SONG, 2015).

Apparently most of the key proteins identified from this study on the up-regulated genes in Myocardial Infarction are more or less connected with cardiac functioning and mostly play a protective role. These findings can be of use when it comes to discovering/designing new drugs in order to limit the damaging effects of myocardial infarction.

## conclusion

This study tried to elucidate the underlying molecular mechanism of myocardial infarction. It identified 11 genes significantly up-regulated in myocardial infarction that corresponded with NCI generated annotations. Five transcription factors, 5 microRNAS and 3 pathways that are significantly associated with them have also been detected. Most importantly, 6 key proteins have been identified from the protein-protein interaction network. These proteins may hold the key to discovering/designing new drugs containing the ability to lower the damages caused by myocardial infarction.

